# Efficient quantification of lipid packing defect sensing by amphipathic peptides; comparing Martini 2 & 3 with CHARMM36

**DOI:** 10.1101/2022.03.04.482978

**Authors:** Niek van Hilten, Kai Steffen Stroh, Herre Jelger Risselada

## Abstract

In biological systems, proteins can be attracted to curved or stretched regions of lipid bilayers by sensing hydrophobic defects in the lipid packing on the membrane surface. Here, we present an efficient end-state free energy calculation method to quantify such sensing in molecular dynamics simulations. We illustrate that lipid packing defect sensing can be defined as the difference in mechanical work required to stretch a membrane with and without a peptide bound to the surface. We also demonstrate that a peptide’s ability to concurrently induce excess leaflet area (tension) and elastic softening – a property we call the ‘characteristic area of sensing’ (CHAOS) – and lipid packing sensing behavior are in fact two sides of the same coin. In essence, defect sensing displays a peptide’s propensity to generate tension. The here-proposed mechanical pathway is equally accurate yet, computationally, about 40 times less costly than the commonly used alchemical pathway (thermodynamic integration), allowing for more feasible free energy calculations in atomistic simulations. This enabled us to directly compare the Martini 2 and 3 coarse-grained and the CHARMM36 atomistic force-fields in terms of relative binding free energies for six representative peptides including the curvature sensor ALPS and two antiviral amphipathic helices (AH). We observed that Martini 3 qualitatively reproduces experimental trends, whilst producing substantially lower (relative) binding free energies and shallower membrane insertion depths compared to atomistic simulations. In contrast, Martini 2 tends to overestimate (relative) binding free energies. Finally, we offer a glimpse into how our end-state based free energy method can enable the inverse design of optimal lipid packing defect sensing peptides when used in conjunction with our recently developed Evolutionary Molecular Dynamics (Evo-MD) method. We argue that these optimized defect sensors – aside from their biomedical and biophysical relevance – can provide valuable targets for the development of lipid force-fields.

**TOC Graphic:** 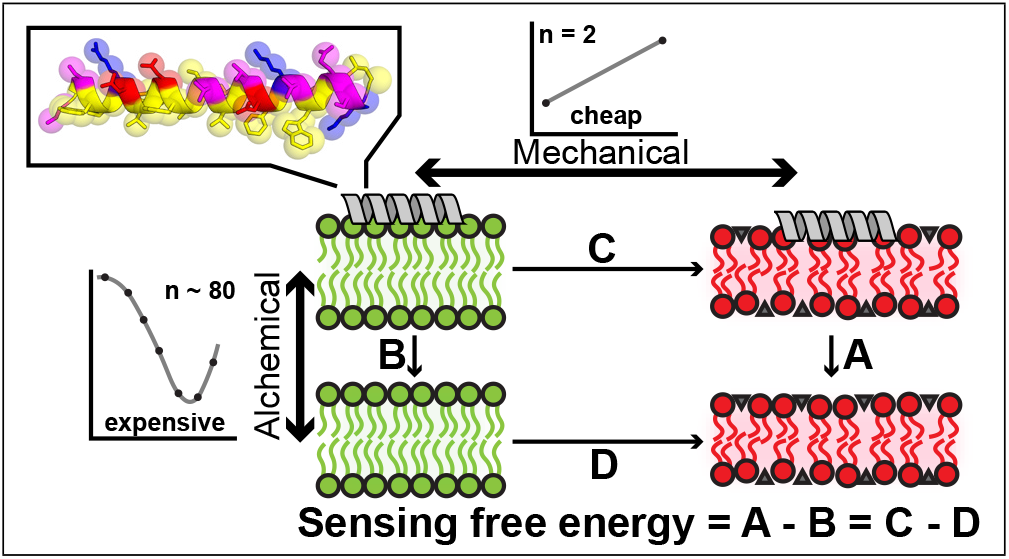

## 1 Introduction

Lipid bilayers are crucial for maintaining cellular integrity, structure and homeostasis. Many processes taking place in, on or near these membranes involve proteins that experience a thermodynamic force that drives the self-organization and recruitment toward certain bilayer properties such as curvature, tension or lipid composition.^1–4^ This process is called ‘sensing’. A key feature that underlies such sensing are lipid packing defects that occur when membranes are stretched or bent. Regardless if this happens symmetrically (i.e. both leaflets experience the same tension) or asymmetrically (i.e. one leaflet gets stretched more than the other, resulting in a net positive curvature), the optimal packing of the hydrophilic lipid head groups gets disturbed, which exposes hydrophobic defects on the membrane surface. Due to the surfactant-like nature of amphipathic peptides and protein motifs, these differences in surface hydrophobicity can give rise to a difference in relative binding free energy (membrane partitioning) and a concomitant sensing force.^5,6^

Many questions related to lipid packing defects can be challenging to address in experiments because these methods lack the required molecular detail. Hence, molecular dynamics (MD) simulations are an indispensable tool to study such membrane properties and their effect on protein binding and sensing. The crucial issue of the reliability of simulations is the quality of the force-field, and many efforts, especially in the latest several years, have been devoted to parametrization and optimization of the force-fields for biomembrane modeling. For example, so called ‘bottom-up’ coarse-grained (CG) models such as the Martini force-field, are parametrized in a systematic way, based on the reproduction of partitioning free energies between polar and apolar phases of a large number of chemical compounds.^7,8^ A main goal of both atomistic and CG lipid force-fields particularly is to accurately reproduce the membrane binding and partitioning of peripheral membrane proteins. However, a systematic comparison of force-fields on the relative binding free energies, i.e. quantification of membrane curvature and lipid packing defect sensing, of whole proteins or even peptides is computationally not tractable using present alchemical approaches such as thermodynamic integration (TI)^9^ and the Bennett acceptance ratio (BAR) method.^10^ Furthermore, (un)binding of peptides is subject to large hysteresis, which complicates accurate estimation of binding free energies when using free energy calculation methods that rely on physical rather than alchemical reaction coordinates (e.g. umbrella sampling). Finally, a prevailing need exists to develop methods that enable efficient and accurate quantification of a peptide’s ability to sense lipid packing defects because of important pharmaceutical applications such as the design of broad-spectrum antiviral peptides that selectively target the highly curved lipid envelope of clinically relevant viruses. ^11^

We recently illustrated how relative binding free energies due to differences in membrane curvature^12^ and lipid packing defects^13^ can be quantified in CG molecular simulations via umbrella sampling. With these studies, we also showed that curvature and lipid packing defect sensing are in fact equivalent phenomena, implying that a protein’s ability to sense positive membrane curvature can be alternatively inferred from its ability to sense packing defects. Lipid packing defect sensing can be efficiently quantified from the magnitude of a thermodynamic sorting force that acts on a surface binding peptide when it is positioned within a spatial gradient of lipid packing defects – a defect gradient (Fig. 1A).^13^ Since the sorting force is approximately constant over the whole gradient (slope in Fig. 1B), its magnitude is directly proportional to the relative free energy of membrane binding. Therefore, sensing can be quantified by ensemble averaging over only a single simulation. This approach yields accurate results given that the spatial gradient zone is smeared out over ≈ 10 nm or more. However, the method still features a slow convergence of the sorting force via ensemble averaging (multiple microseconds), due to the asymmetric nature of the gradient in combination with slow orientational and rotational modes of the peptide when it is bound to the membrane. Furthermore, computational efficiency can not be trivially improved via further reduction of system size, since too strong spatial gradients in membrane thickness also compromise the precision of the method. Thus, although the thermodynamic gradient method provides an elegant and intuitive method for quantifying lipid packing defect sensing, its slow convergence limits its application in high-throughput utilities and atomistic simulations.

**Figure 1:**
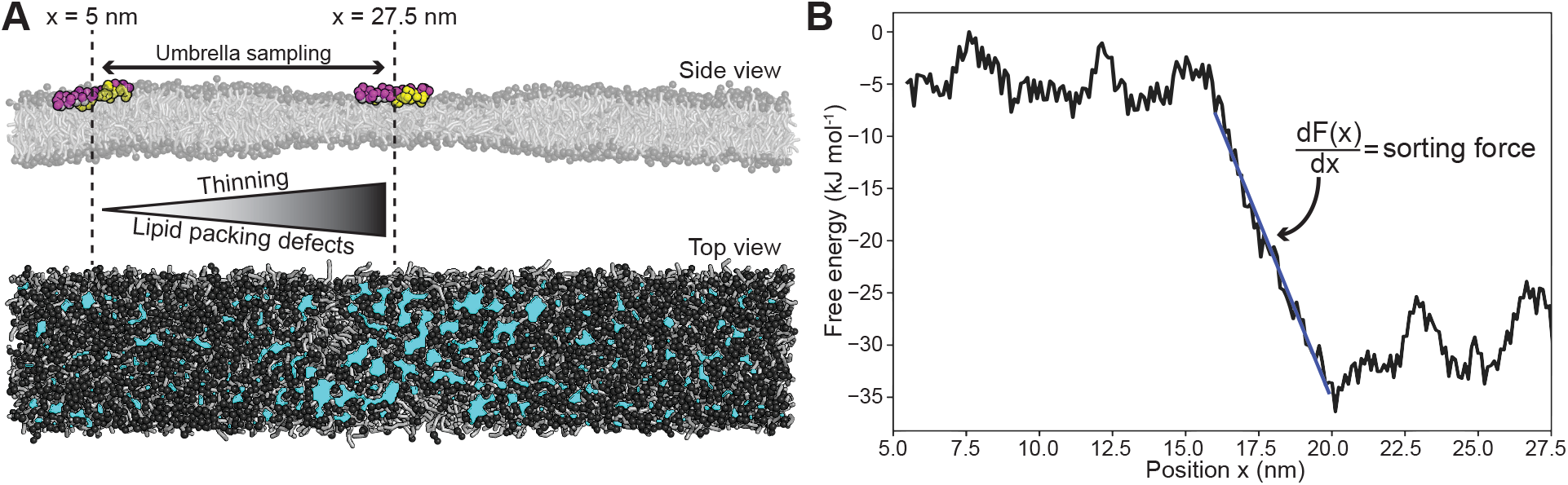
Qauntification of lipid packing defect sensing using the thermodynamic gradient method. The figure is adapted from our previous work.^13^ A) Side view and top view of a flat Martini POPC membrane subject to an external potential that induces tension (thinning) in the middle section (*x* = 27.5 nm, thickness ≈ 3 nm) and is gradually switched off moving outward (to *x* = 5 nm, thickness ≈ 4 nm). The peptide model is ALPS. Cyan patches in the top view indicate lipid packing defects.^14^ B) The free energy (*F*(*x*)) as a function of the position (*x*) on the membrane calculated by umbrella sampling for ALPS across the lipid packing defect gradient. The slope of this curve 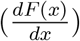 is a thermodynamic sorting force, which can be directly used as a measure for lipid packing sensing. The linear behavior is explained by the linear decrease in membrane thickness along the gradient. Owing to bulk incompressibilty and hookian membrane elasticity, this results in a linear gradient in surface tension and surface free energy whose spatial derivative yields a constant sorting force along the gradient.

Here, we will present a highly efficient and accurate end-state free energy calculation method to directly estimate the relative free energy of membrane binding, i.e. the quantification of defect and curvature sensing. In contrast to well-established *alchemical pathways*, we will illustrate that the relative free energy of binding can be alternatively obtained using a *mechanical pathway*, which is equally accurate and computationally much less expensive. An important advantage of the new approach is that the corresponding end-state systems are both small (128 lipids) and symmetric, which enables quick and reliable calculation of the relative binding free energy in CG simulations and also makes them more feasible to conduct on the atomistic scale. We will compare the performance and accuracy of this endstate mechanical pathway with the standard alchemical pathway (TI) by analyzing them for four known packing defect sensing peptides and two related negative controls. We will also use this new quantification method to compare lipid packing defect sensing and overall peptide-membrane interactions within the recent Martini 3 force-field^8^ with the previous version and atomistic simulations.

Importantly, the here-resolved thermodynamic cycle illustrates that lipid packing sensing and the induction of membrane tension are in fact two sides of the same coin. This implies that defect sensing peptides maximize the generation of leaflet tension resulting in a strong native membrane destabilizing propensity. Finally, as the ultimate demonstration of the method’s high-throughput potential, we will illustrate how our method can direct the simulated evolution of defect sensing peptides toward optimal sensing (inverse design). We argue that these optimal sequences provide novel and valuable benchmark systems since they reflect the boundary of a force-field’s applicability domain. The primary aim of force-fields is to reproduce physicochemical driving forces – trends, not absolutes – and, therefore, they must at least reproduce the global physicochemical features of optimized sequences.

## 2 Theory and methods

### 2.1 Alchemical calculation of the relative binding free energy

Lipid packing defect sensing can be defined as the differential affinity of a peptide toward a membrane *with* defects (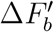, under tension) versus a membrane *without* defects (Δ*F_b_*, tensionless). Note that we use the nomenclature Δ*F* to indicate that the simulations were performed at a constant area (but not constant volume). In terms of free energy differences, we can write:

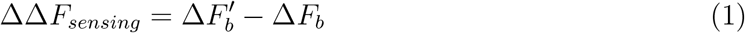

These terms can be calculated using thermodynamic integration (TI). ^9^ In this commonly used ‘alchemical’ method, a peptide-membrane bound state (A, with potential energy *V_A_*) is transitioned to the peptide *in vacuo* (B, with potential energy *V_B_*) by gradually switching off the Van der Waals- and Coulomb-interactions between the peptide particles and their surrounding (coupling parameter λ = 0 → λ =1). By taking the integral of the ensemble average of the derivative of the potential energy (*V*(λ)), the free energy difference between state A and B can be calculated:

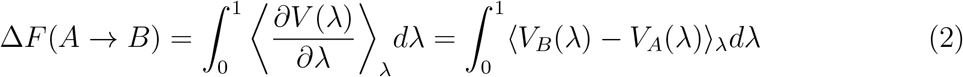

Although TI is an accurate and commonly used method, it is – even when using CG forcefields – computationally expensive. To make sure the numerical integration is valid, a smooth 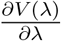-profile is required, which typically takes at least 20 λ-states to simulate. Because decoupling is generally performed separately for the Van der Waals- and Coulomb-interactions, the number of simulations increases to 30-40 per system. Moreover, in problems concerning the difference in binding free energies between two systems (like our packing defect sensing problem; 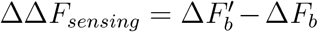), this again doubles to at least 60-80 simulations in total.

### 2.2 Mechanical calculation of the relative binding free energy

Consistent with the first law of thermodynamics, the energy difference between two states of a cycle is independent of the path one takes to get from one to the other. We can define such a cycle by connecting the two begin-states (peptide bound) and the two end-states of the alchemical pathway (peptide unbound) – see Fig. 2A. This realization allows a redefinition of lipid packing defect sensing as the change in work required to stretch a membrane in presence 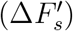 and absence (Δ*F_s_*) of a surface-bound peptide:

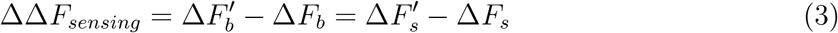

**Figure 2:**
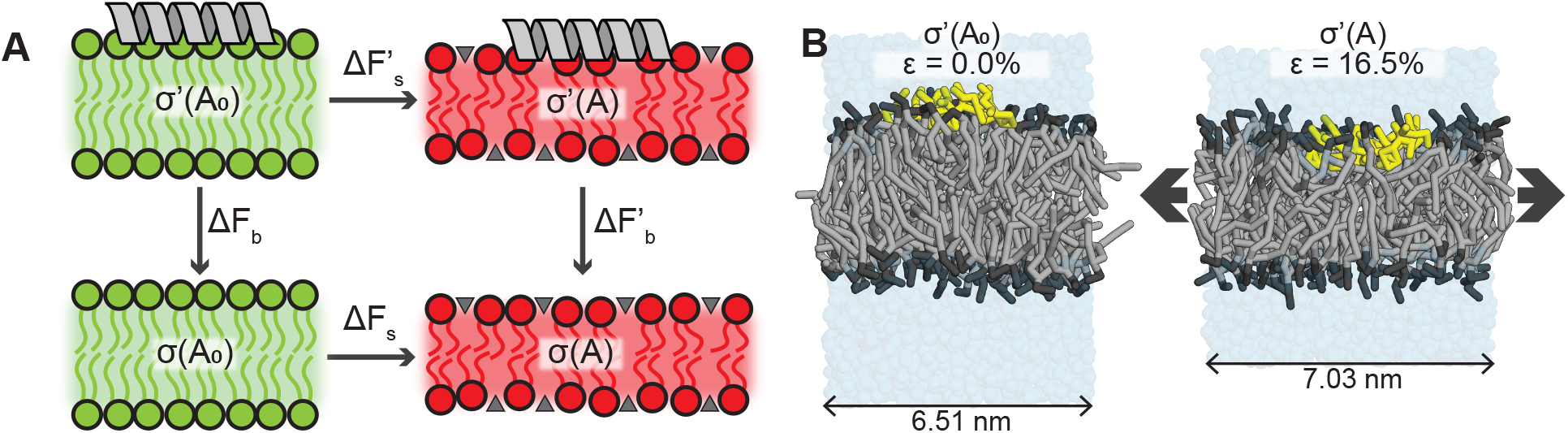
A) Thermodynamic cycle that links the alchemical 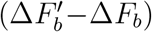 and the mechanical 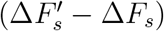 pathways. Tensionless and stretched membranes are shown in green and red, respectively. Lipid packing defects in the stretched membranes are depicted by the gray triangles. B) Snapshots of the CG systems without tension (left, no relative strain *ϵ*) and with tension (right, high relative strain *ϵ*). POPC lipids are shown in gray, with black head groups. The peptide is shown in yellow.

Thus, a good lipid packing sensor (ΔΔ*F_sensing_* << 0) minimizes the work required to stretch the membrane leaflet it adheres to.

Calculating this mechanical pathway is much cheaper compared to the alchemical pathway (TI) for two reasons. First, one of the terms, Δ*F_s_*, is peptide-independent, since it is simply the work required to stretch a membrane without a peptide bound to it. This means one only has to calculate it once (for a given system) and can simply plug in the same number for any peptide of interest. Second, in elastic theory, the lateral tension *σ*(*A*) in a membrane is linearly related to the change in membrane area (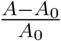, i.e. relative leaflet strain *ϵ*) for small deviations from the equilibrium tensionless area *A*_0_;^15^

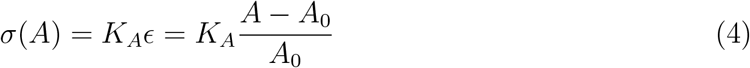

with *K_A_* being the area compressibility modulus. The tension *σ*(*A*) can be directly obtained via ensemble averaging at constant membrane area using the relation

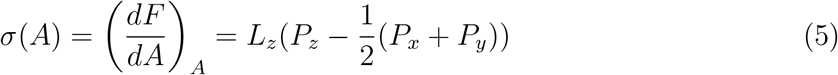

where *L_z_* is the length of the simulation box along the *z*-dimension and *P_x_*, *P_y_*, and *P_z_* are the average pressures in the *x*-, *y*-, and *z*-dimensions, respectively. These pressures are derived from the diagonal components of the stress tensor as derived from the Clausius virial theorem.

Because of the linear relation in eq. 4, the free energy difference (mechanical work) of stretching (Δ*F_s_*) can be reliably approximated by performing MD simulations at merely two different, constant areas *A* and *A*_0_ and by measuring the resulting surface tensions *σ*(*A*) and *σ*(*A*_0_). Applying the trapezoidal rule yields:

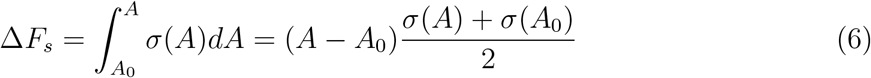

This is where the biggest efficiency gain lies for the mechanical path (two points on a straight line) versus the alchemical path (integration over >20 points on a complicated 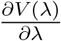-profile).

Now, combining and rearranging eq. 3 and eq. 6 and yields:

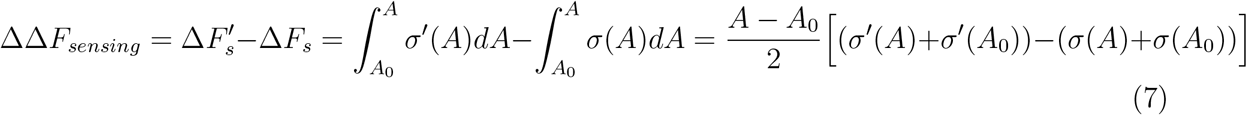

In which – like mentioned before – the membrane tensions in the peptide-free systems *σ*(*A*) and *σ*(*A*_0_) only have to be measured once to be used in the calculations for any peptide.

### 2.3 System set-up

A simulation box containing 128 CG 16:0–18:1 phosphatidylcholine (POPC) lipids was generated using the *insane* python script. ^16^ After solvation with Martini water beads and a steepest descent minimization, the system was equilibrated with semiisotropic pressure coupling to obtain tensionless membrane conditions. Next, the area of the box was increased by steps of 1.4 nm^2^ and reequilibrated every time (at constant area). 500 ns production runs were performed for the resulting 6 membranes, the areas of which ranged from 42.4 nm^2^ to 49.4 nm^2^ (Fig. 2B). Such a 16.5% increase in leaflet area corresponds to the effective relative strain 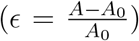 in the outer leaflet of a ≈ 25 nm diameter vesicle (see SI), which is the lower size limit of vesicles found in nature.

Helical CG peptide models were generated using PeptideBuilder^17^ in conjunction with martinize2 – VerMoUTH^18^ and placed 1.5 nm from the membrane center plane. A steepest descent minimization with soft-core potentials (0.75 coupling for the Van der Waals interactions) was performed to solve clashes. Production runs of 1-5 *μ*s were performed, the first 50 ns of which were discarded from further analyses for equilibration.

For TI, the equilibrated setups for the minimal and maximal tension membranes were used as the starting points. 37 λ-states were defined to ensure proper 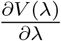-sampling during separate decoupling of the Van der Waals- and Coulomb interactions (37 × 2 = 74) to the final in *vacuo* state. For the two setups – low and high tension – a total of 74 × 2 = 148 simulations of 500 ns was performed, for each peptide. The Langevin stochastic dynamics (SD) integrator and thermostat were used for these runs. A soft harmonic distance constraint (*k_force_* = 50 kJ mol^−1^ nm^−2^) was used between the centers of mass of the membrane and the peptide (*z*-dimension only) to prevent peptide-membrane dissociation in high decoupling states. Finally, free energy differences were obtained through numerical integration (eq. 2).

For buckled membrane simulations, initial configurations were generated with the python script *insane*.^16^ Each membrane leaflet is comprised of 767 POPC lipids, which were put in the *x-y* plane of a 40 nm · 10 nm · 20 nm simulation box. A curvature sensing peptide was placed close to the upper leaflet. The system was solvated with standard Martini water and a 0.15M NaCl concentration. Following steepest-descent energy minimization and initial equilibration (50 ns *N_p_T*) the system was compressed in x-direction by applying a pressure of 3 bar (Berendsen barostat,^19^ *τ_p_* = 12.0 ps, compressibility of 3 · 10^−4^ bar^−1^). The system was allowed to expand in *z*-direction, while the *y*-dimension was fixed. From the compression trajectory, a frame close to the compression of 38% was chosen. Another *N_p_T* equilibration with fixed *x*- and *y*-dimensions (100ns, Parrinello-Rahman barostat,^20^ *τ_p_* = 12.0 ps, compressibility of 3 · 10^−4^ bar^−1^) followed. To fix the analytical shape of the buckled membrane positions of the PO4 beads in the lower leaflet were restrained with a force constant of 10 kJ mol^−1^ nm^−2^. This minimally influences the upper leaflet dynamics, where the peptide is located. For subsequent peptides the preequilibrated system was used, the original peptide was deleted and a new peptide was inserted into the free volume. Energy minimization and *NVT* equilibration followed. To generate initial configurations for the umbrella sampling, the peptide was pulled along the x-direction of the membrane. Each individual umbrella sampling run is 1.05 *μ*s long, with the first 50 ns for equilibration. A harmonic potential with a force constant of *k* = 100 kJ mol^−1^ nm^−2^ was used to restrain the peptide to a defined point along the reaction coordinate. Rotation around the membrane normal was restrained in the same manner to keep the peptide aligned with the y-axis, i.e. the flat direction of the membrane.

Atomistic POPC membranes were equilibrated at the same constant areas as in the CG simulations. The helical atomistic peptide models were built by feeding the PeptideBuilder pdb files into the CHARMM-GUI.^21^ The resulting peptide structures were placed on the membranes (preequilibrated in tip3p water) following the same procedure as described for the CG simulations. Production runs of 1 *μ*s were performed. The first 500 ns were not included in the analyses to ensure proper equilibration.

### 2.4 Simulations details

All simulations were performed with GROMACS 2019.3,^22^ except for the simulations on a buckled membrane, which were done with GROMACS 2021.4. The temperature was kept at a constant 310 K by the velocity rescaling thermostat^23^ (*τ_T_* = 1 ps). Unless stated otherwise, simulations were performed with fixed *x*- and *y*-dimensions (constant area), i.e. Berendsen pressure coupling^19^ was applied only in the *z*-dimension (1 bar reference pressure with a 4.5 · 10^−5^ bar^−1^ compressibility).

Coarse-grained molecular dynamics (CGMD) simulations were performed with the Martini force-field, version 3.0.0,^8^ unless stated otherwise. A 30 fs time step was used for all CG simulations except the buckled membrane runs, which were performed with a 20 fs time step. Van der Waals interactions were calculated with the shifted Verlet cut-off scheme^24^ and reaction-field electrostatics^25^ describe the coulomb potentials, both with a 1.1 nm cut-off. The neighbor list was updated every 20 steps.

Atomistic simulations were done with the February 2021 version of the CHARMM36 force-field.^26,27^ A time step of 2 fs was used. Van der Waals and Coulomb interactions were calculated using the shifted Verlet^24^ and particle mesh Ewald (PME)^28^ methods, respectively, both with a 1.2 nm cut-off distance. The neighbor list was updated every 10 steps. The LINCS algorithm^29^ was used to constrain bonds with hydrogen atoms.

## 3 Results and discussion

To demonstrate how our method works and to validate it against the well-established TI method, we will focus on the most broadly studied class of lipid packing sensing protein motifs: amphipathic *α*-helices. One side of these peptides mainly consists of large apolar/aromatic moieties to complement the hydrophobic lipid packing defects and the other side comprises polar and/or positively charged residues to interact with the solvent and lipid headgroups. We picked six peptides for our study (Fig. 3A), which we will briefly introduce below (see Table SI1 for details).

**Figure 3:**
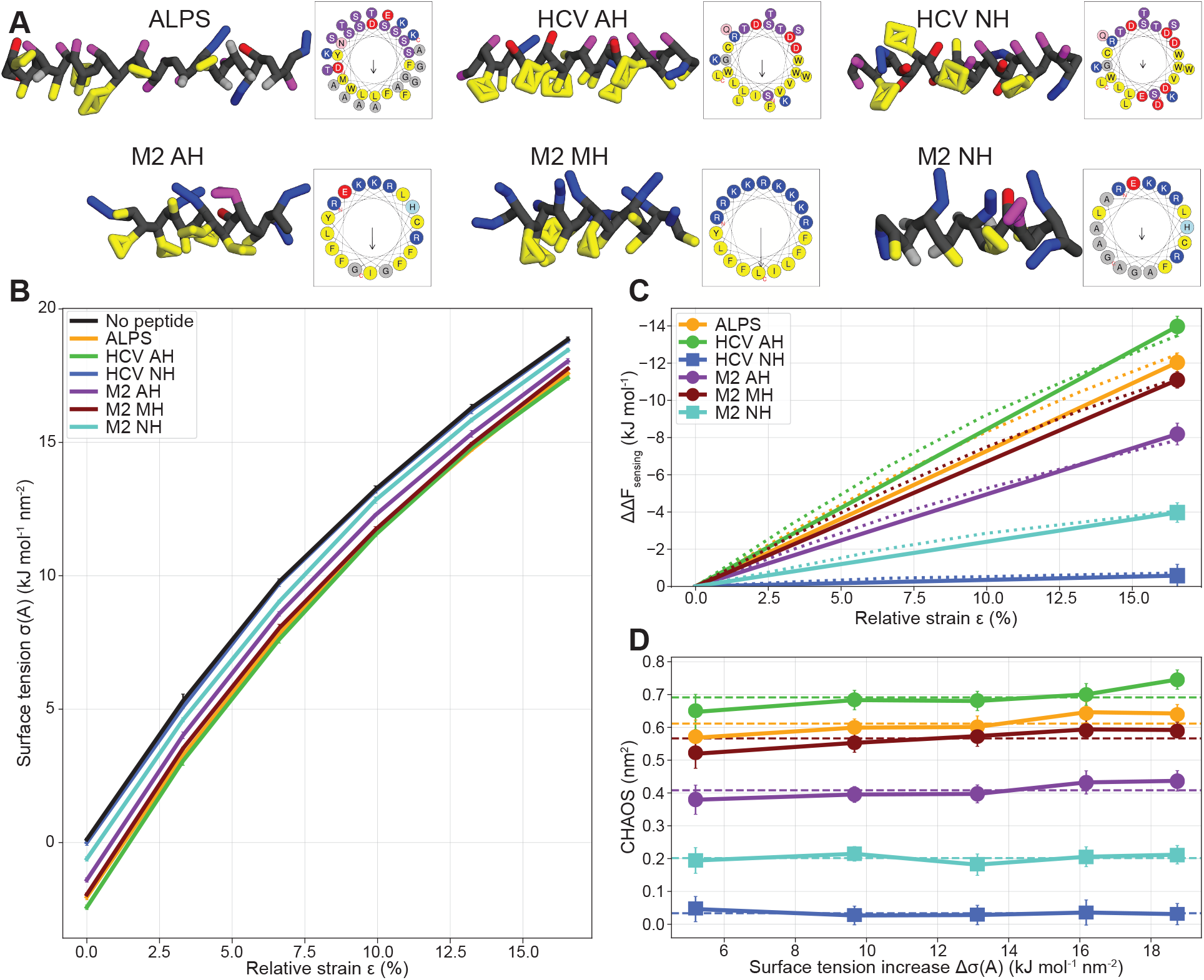
A) Depictions of the helical peptide models in the Martini 3 force-field,^8^ and full-sequence helical wheel representations.^42^ Yellow: hydrophobic residues (C, F, I, L, M, V, W, Y). Red: negatively charged residues (D, E). Blue: positively charged residues (K, R). Magenta: polar residues (H, N, Q, S, T). Gray: small residues (A, G) and backbone. B) The surface tension measured at increasing membrane areas (relative strain 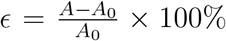) with and without surface-bound peptides. C) Sensing free energy differences at increasing relative strain, as calculated with eq. 7, for different peptides. The dotted line represents the integral over all six points. The solid line represents the end-state method, which only integrates between the first (*ϵ* = 0%) and the last (*ϵ* = 16.5%) point. D) The characteristic area of sensing (CHAOS parameter, see eq. 8) is constant for different relative surface tension values (Δ*σ*(*A*) = *σ*(*A*_0_) – *σ*(*A*)) thereby illustrating its end-state invariant nature. The dashed lines represent the average values for each peptide.

First, we will study the ALPS motif that allows curvature sensing by the ArfGAP1 protein,^30^ and is also found on other proteins.^31–34^ Since its discovery, ALPS has served as an important model peptide in many computational studies on curvature/lipid packing sensing,^35–37^ also in our own group.^12,13^ Second, we will include an amphipathic helix (AH) that was derived from the NS5A protein of hepatitis C virus (HCV) and discovered to sense and rupture vesicles in a size-dependent manner: small vesicles (including HCV particles themselves) were more readily ruptured than bigger ones and this size range overlaps with the diameter of many enveloped viruses (50-160 nm).^38,39^ And, indeed, antiviral activity of HCV AH was later found for several unrelated viruses, including Zika, Dengue and West Nile virus. ^40^ Just like the original work, we will also include the negative control HCV NH, where three point mutations nullify the amphipathicity of the peptide, resulting in a loss of antiviral activity. ^38^ Finally, and along the same lines, we will consider an AH derived from the M2 protein of the influenza virus that showed antiviral activity against four different influenza strains.^41^ A variant with increased amphipathicity (named M2 MH) showed an 16.4-fold increase in anti-influenza inhibitory effect. In contrast, the potency is abolished for a variant with low amphipathicity (named M2 NH), which we will use as a second negative control.

### 3.1 Calculating sensing free energies via the mechanical pathway

We performed CGMD simulations of these peptides adhered to the surface of membranes at equilibrium (0% relative strain) and at increasing degrees of stretching (up to 16.5% relative strain) and measured the resulting change in surface tension *σ*(*A*) (Fig. 3B). Consistent with eq. 4, we observed near-linear behavior for small strains both with and without peptides present. It becomes clear that peptides binding to the surface of a membrane reduce the surface tension imposed by the fixed boundary conditions to keep the system at a constant area. In other words, the adhered peptides reduce the work of stretching. Also, we can already observe that the inactive peptides HCV NH and M2 NH cause a much smaller reduction in this tension than the active curvature sensors. Now, we can calculate the free energy of sensing (ΔΔ*F_sensing_*) by integrating over these curves (eq. 7). I.e. we calculate the area enclosed by the ‘no peptide’ curve (*σ*(*A*); black line in Fig. 3B) and the curve for the peptide of interest (*σ*′(*A*); colored line). Since the tension reduction is approximately constant for different membrane areas, taking this integral over all 6 points (dotted line in Fig. 3C) or only the end-states (0% and 16.5% relative strain; solid line in Fig. 3C) yields the same result, or at least within the measurement error. Thus, only two simulations at the extremes suffice to accurately calculate ΔΔ*F_sensing_*. We should note that this ΔΔ*F_sensing_* is calculated from simulations at constant area. With additional simulations at constant tension (see SI), we show that the resulting correction terms that arise from transitioning from constant area to constant tension ensembles are negligible. Therefore, we proceeded to use the endstate mechanical calculations at constant area for all following results described in this paper.

### 3.2 The ‘CHAOS’ parameter: an end-state invariant measure for lipid packing sensing ability

We should note that one can only interpret the ΔΔ*F_sensing_* relative between different peptides, since the absolute values depend on the (arbitrary) choice of the two end-states (Fig. 3C). We chose end-states with a large difference in tension, since this inflates the value of ΔΔ*F_sensing_*, and therefore enhances the reliability of peptide ranking, i.e. the relative differences in ΔΔ*F_sensing_* overcome the sampling error. In fact, because of a linear relationship between free energy and tension (eq. 6), we can define an end-state invariant property that we coin the ‘characteristic area of sensing’ (aptly abbreviated to ‘CHAOS’), since it has the dimension of area per molecule. This CHAOS parameter is the relative difference in binding free energy (=sensing) normalized by the difference in surface tension between the peptide-free reference systems:

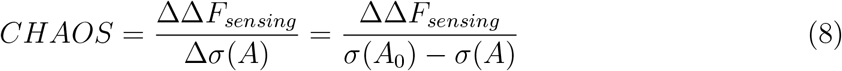

The surface tension of the peptide-free reference systems is chosen pragmatically because of the smaller sampling error in pure lipid membrane systems. In the constant tension ensemble both systems in fact converge to the same surface tension, which yields similar magnitudes of the CHAOS parameter (see SI). Indeed, when we plot the CHAOS parameter against the range of Δ*σ*’s simulated, we observe an approximately constant CHAOS value for every peptide (Fig. 3D). The ranking and relative distances between these values for the different peptides remains fully consistent with the ΔΔ*F_sensing_* values, yet they are independent of the choice of end-states.

The CHAOS parameter can be, in part, interpreted as the area increase upon adhesion of a given peptide to a tensionless membrane within the corresponding *N_p_T* ensemble (see Table SI2). However, its magnitude will always be slightly bigger, since it is additionally determined by the peptide’s ability to soften the membrane, i.e. lowering the area compressibilty *K_A_*, and thereby reducing mechanical work of stretching. In fact, ‘CHAOS’ is a striking abbreviation since its magnitude directly reflects the ability of the peptide to disorder the packing of lipid tails via the creation of excess leaflet area (tension). As per this definition, we speculate that CHAOS parameters may be directly comparable to experimental values obtained from Langmuir-Blodgett experiments in which the surface-tension of a lipid monolayer at the air-water interface can be measured upon adhesion of surfactants (e.g. amphipathic peptides).^43^

### 3.3 Convergence, reproducibility and comparison to conventional alchemical pathways

We performed three independent reruns of 5 *μ*s per simulation to assess the reproducibility and convergence of our method (Fig. 4A). This showed that after approximately 1 *μ*s, the calculated free energies of the independent runs converged to the same value (within the margin of error).

**Figure 4:**
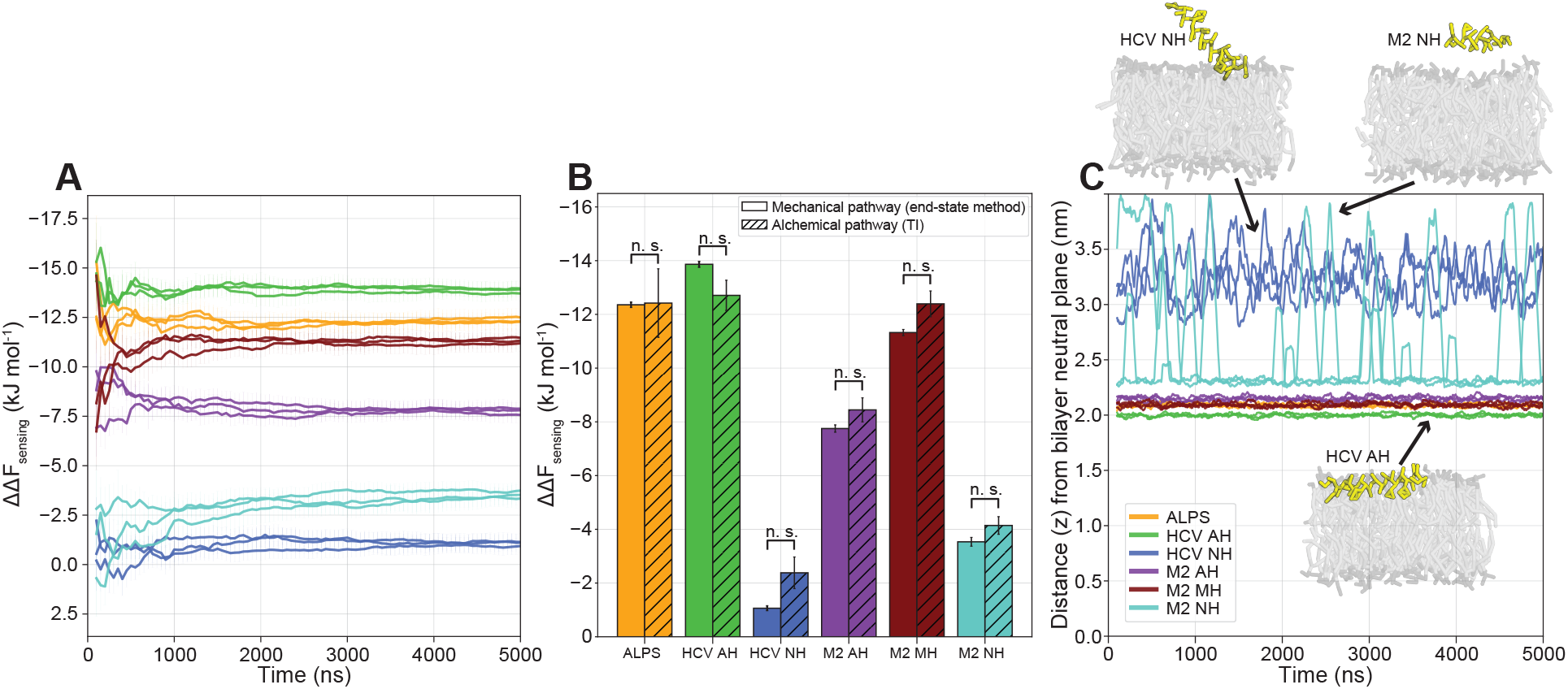
A) Cumulative moving average of ΔΔ*F_sensing_* for triplicate 5 *μ*s runs. B) Comparison between our mechanical end-state method (average of the three 5 *μ*s runs in Fig. 4A) and free energy calculation via the alchemical pathway (TI). P-values were calculated with the two-tailed Welch’s t-test, which showed no statistically significant differences for any of the peptides (*p* > 0.05). C) The distance (*z*) between the peptide and the bilayer neutral plane (100 ns moving average) for the tensionless runs in Fig. 4A. Absolute values are taken to account for periodic boundary crossings. Insets show typical conformations for HCV AH (fully membrane bound), HCV NH (partly membrane bound) and M2 NH (fully unbound).

Consistent with thermodynamic theory (eq. 3, Fig. 2A), the sensing free energies calculated via the mechanical pathway between the end-states closely match the free energies calculated by TI (Fig. 4B, Fig. SI2), with no statistically significant differences between the two methods for all peptides tested. We would like to stress that, whilst producing the same results, there is a significant computational speed-up for our mechanical end-state method compared to TI. Free energy calculation via the mechanical pathway can be calculated reliably in 2 *μ*s of simulation (1 *μ*s for each of the two states), whereas TI required 74 μs. This 37x speed-up is indispensable when considering high-throughput calculation of membrane binding properties of peptides.

In line with the experimental results and the original design principles, the peptides with mutated hydrophobic faces (HCV NH and M2 NH) indeed have a compromised sensing ability compared to the original peptides that were derived from (HCV AH and M2 AH, respectively; see Fig. 4B). This reduced sensing free energy is mainly due to (partial) detachment from the membrane during our MD simulations (Fig. 4C), which renders them incapable of reducing the membrane surface tension.

### 3.4 Transferability between tension sensing and curvature sensing

In order to evaluate the transferability between lipid packing defect sensing in a flat membrane (tension in both leaflets) and positive curvature (tension in the outer, compression in the inner leaflet), we performed umbrella sampling (US) along a buckled POPC membrane (Fig. 5A-B), as described previously. ^12^ The poorly binding HCV NH and M2 NH peptides were not considered here, since surface adhesion is a prerequisite in this approach. From this experiment, we obtained sensing free energy profiles as a function of membrane curvature (Fig. 5C), that we can directly compare with the free energy values we obtained through our end-state mechanical method (Fig. 3C). We find that US along the membrane buckle yields a similar ranking as our end-state free energy calculation method with flat membranes, with the exception that ALPS outperforms HCV AH by 0.65 kJ mol^−1^ on the buckle (Fig. 5C). However, considering the large error in the US method (95% confidence intervals: 1.57 and 1.34 kJ mol^−1^, respectively), we do not consider this difference to be significant. Thus, the ability of an *α*-helical peptide to sense leaflet curvature is directly relatable to its ability to sense lipid packing defects in a flat membrane.

**Figure 5:**
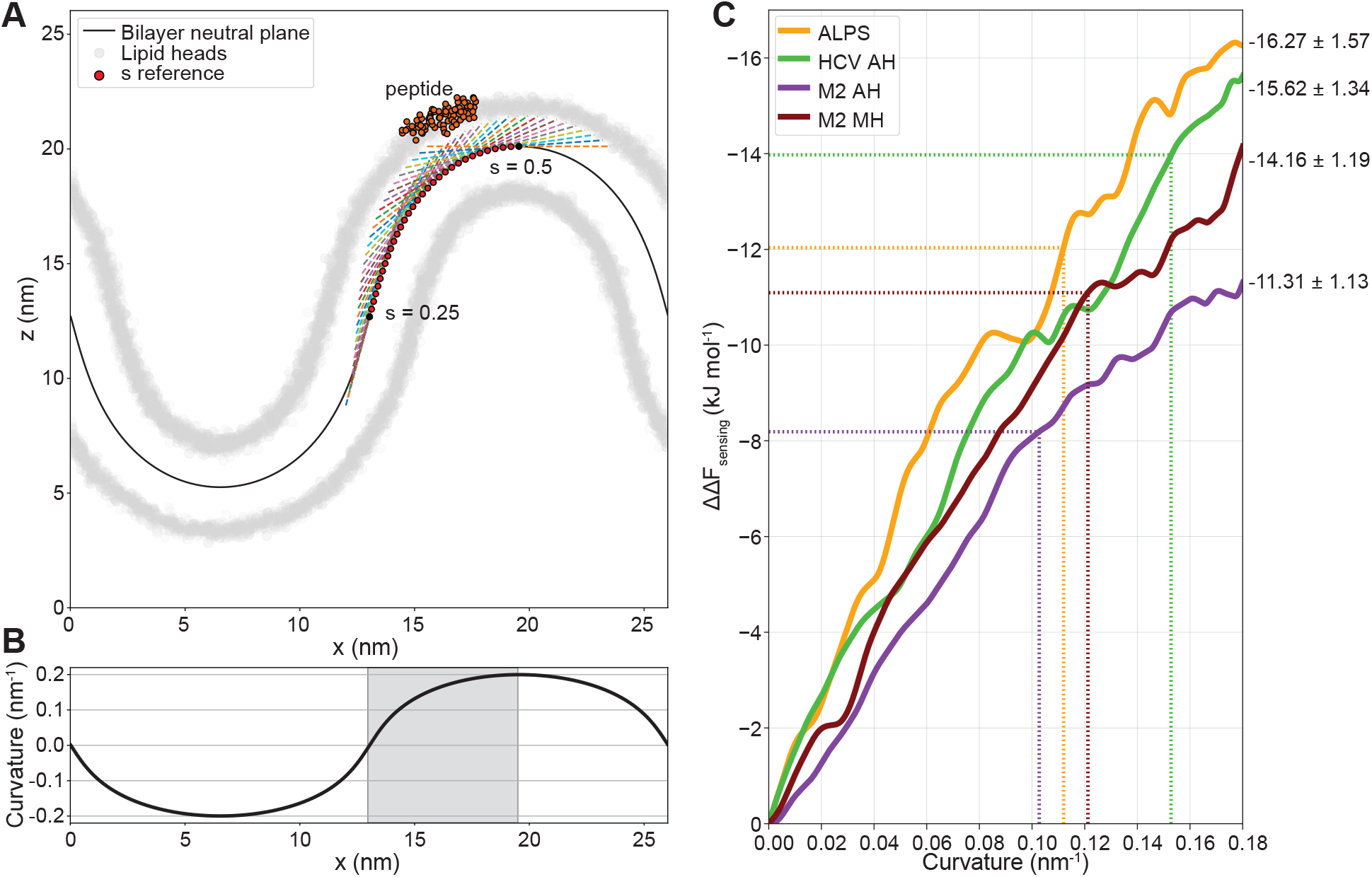
A) Setup for umbrella sampling of peptide-membrane interaction along a buckled membrane. The umbrella potential acts along the tangent (dashed lines) at select reference points (*s*). B) Curvature of the buckled bilayer neutral plane as a function of the *x*-coordinate. The gray area indicates the sampled region. C) Sensing free energy as a function of curvature along the buckled membrane. Dotted lines indicate the free energies values calculated from the end-state mechanical method (Fig. 3C) and their corresponding curvatures.

Furthermore, we can use this membrane buckle to check whether the degree of stretching we use in our end-state method realistically represents the membrane curvature that it should resemble (1/*R* ≈ 1/12.5 = 0.08 nm^−1^, see derivation in SI). We do this by matching the ΔΔ*F_sensing_* values we obtained from our end-state method with the free energy profiles from US over the buckled membrane (Fig. 5C), which yields curvatures ranging from 0.11 to 0.16 nm^−1^ (dotted lines in Fig. 5C). Since the buckled membrane has a cylindrical geometry (only curved in one dimension), the mean curvatures of a corresponding vesicle (curved in two dimensions) should be reduced by a factor two: 0.055 to 0.080 nm^−1^. This results in a range of vesicle radii of approximately 12.5 to 18 nm, which is in line with the estimated vesicle sizes in the tensed membrane systems based on leaflet strain elastic theory as derived in the SI.

### 3.5 Comparing lipid force-fields on sensing free energy and CHAOS parameters

A major goal in the parametrization of atomistic and CG lipid force-fields is to reproduce the partitioning free energies of biomolecules between polar (solvent) and apolar (lipid membrane) phases. Recently, it was shown that the new Martini 3 model is able to correctly characterize the general binding behavior of membrane peripheral proteins.^44^ In addition, we now have a unique tool in hand to quantitatively compare force-fields on their ability to reproduce thermodynamic properties associated with membrane peripheral protein binding such as relative binding free energies (lipid packing sensing) and the concomitant CHAOS parameter. Here, we perform such a comparison for Martini 3, Martini 2 (version 2.2^7,45^ and version 2.3P with polarizable water (PW)^46^ and PME electrostatics) and the atomistic CHARMM36 force-field (Fig. 6A-B). For the six peptides we study throughout this paper, we find that the new Martini 3 model qualitatively reproduces the general trends derived from experimental studies: HCV AH outperforms the inactivated mutant HCV NH^38^ and M2 MH is more potent than M2 AH, whilst the activity is indeed strongly reduced for M2 NH. ^41^

**Figure 6:**
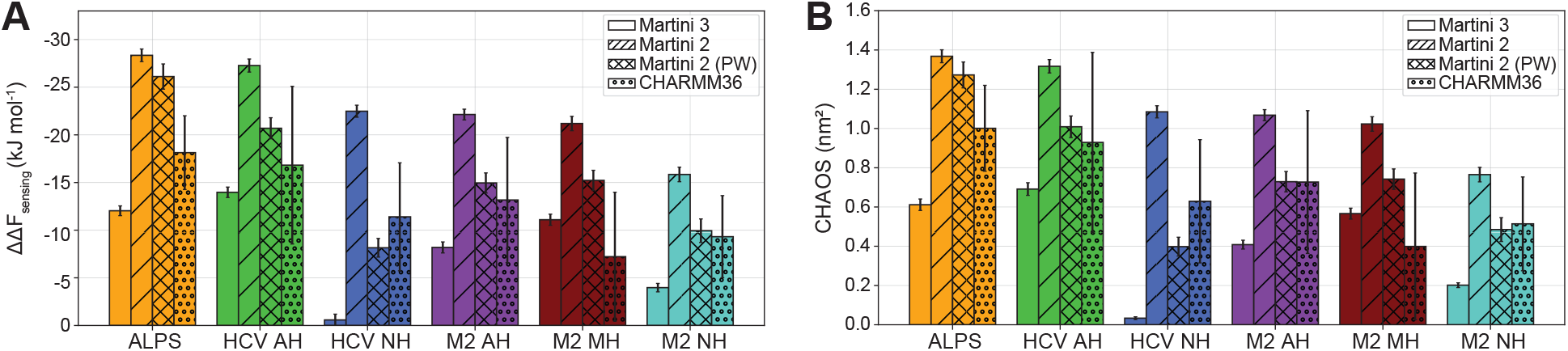
Comparison of lipid packing sensing of peptides with the Martini 3 (version 3.0.0), Martini 2 (version 2.2), Martini 2 with polarizable water (PW, version 2.3P) and atomistic CHARMM36 force-fields in terms of sensing free energy (A) and CHAOS parameter (B). Error bars represent standard errors estimated by five-fold block averaging over the trajectories.

In contrast, these trends are not captured correctly by the Martini 2 model, which severely overestimates ΔΔ*F_sensing_* and CHAOS values compared to the other force-fields. This is caused by exaggerated peptide-membrane binding (i.e. the peptides are ‘too hydrophobic’), as confirmed by the density plots and insertions depths (see Fig. SI3 and Table SI3). More accurate results are obtained when using Martini 2 with PW, especially for peptides with net charge (e.g. HCV NH and M2 AH), for which it more closely matches the values we obtained with Martini 3 and CHARMM36.

The ΔΔ*F_sensing_* and CHAOS values calculated from atomistic simulations with CHARMM36 are within the same range as the Martini 3 and Martini 2 (with PW) results. However, we should note that measurement errors are considerably larger because of the slower convergence compared to the CG force-fields, rendering sufficient sampling of (un)binding and peptide refolding events challenging in practice despite the high computational efficiency of our method.

Finally, we examined the membrane insertion depths of the peptides in the different forcefields and found that this insertion is markedly shallower in Martini 3 than in CHARMM36, i.e. peptides in Martini 3 behave ‘too hydrophilic’ (see Fig. SI3 and Table SI3) despite qualitatively reproducing the experimental trends. Part of this discrepancy could be related to the structural plasticity of peptides in the atomistic simulation, which makes a direct comparison to CG simulations less straightforward. Nevertheless, the observed systematic and marked reduction in peptide insertion depths along with an often lower CHAOS parameter suggests that Martini 3 may have a tendency to underestimate the membrane binding behavior of proteins with respect to CHARMM36 even though overall relative binding free energies seem improved with respect to the previous Martini versions. One of the main founding principles of the original Martini model was to reproduce partitioning free energies of fluid mixtures by reproducing density profiles, which determine the insertion depths of molecules. It is thus somewhat surprising that the membrane insertion depth of proteins are in fact less well reproduced in Martini 3 than in the older versions.

### 3.6 Using ‘CHAOS control’ to improve force-field development?

In this work, we selected six peptides based on the fact that they are of biological/pharmaceutical importance and experimentally well-characterized. However, the similarity in their CHAOS parameters suggests that peptides can have overlapping physicochemical properties, despite having very different sequences. This raises the question if a physicochemically more diverse set of peptides can be constructed that more efficiently and strategically enables both the benchmarking and parametrization of force-fields.

We recently proposed a physics-based inverse design approach coined Evolutionary Molecular Dynamics (Evo-MD). ^47^ Evo-MD relies on the principle that (large) experimental data contributes to solving biophysical problems independently via the parametrization of bottom-up CG force-fields. Evo-MD features a directed evolution (genetic algorithm) of peptide sequences that starts with a set (‘population’) of completely random peptide sequences (‘individuals’) and computes the corresponding ‘fitness’ value for every sequence from one or more MD simulations. Then, in each generation/iteration, analogous to Darwin’s ‘reproduction of the fittest’ the best scoring individuals are recombined and consequently produce offspring that resemble their predecessors, but are slightly different due to both random cross-overs and infrequent point mutations. By repeating this cycle (Fig. 7A) for several iterations, the fitness of the best individuals in the population should improve until it converges to an optimum value. We have previously demonstrated the utility of this concept by resolving the optimal cholesterol sensing transmembrane domain. ^47^

**Figure 7:**
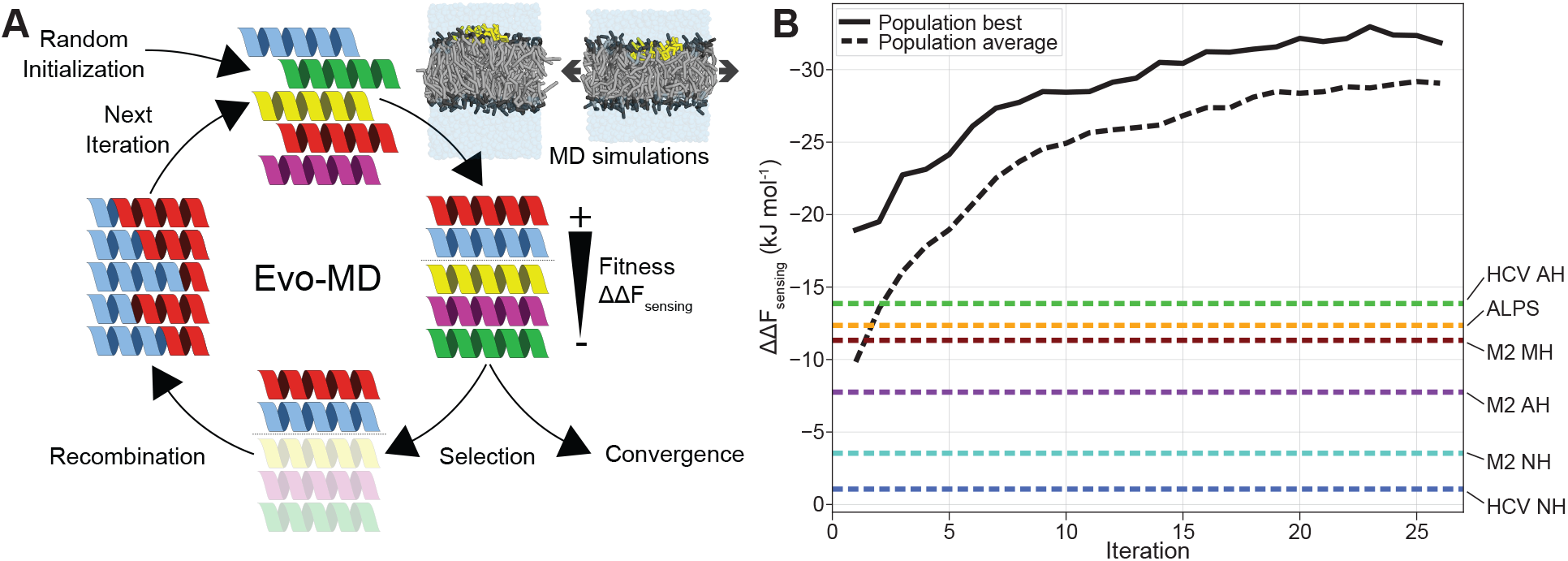
A) Schematic representation of the basic concept of evolutionary molecular dynamics (Evo-MD), adapted from. ^47^ Generated peptides (starting from a population of random sequences) are iteratively ranked on their ‘fitness’ (ΔΔ*F_sensing_*), as determined by the endstate mechanical pathway method described in this paper. The best sequences are picked and recombined to produce the next generation, leading to gradual evolution toward the optimal lipid packing sensing peptides. B) Within 25 iterations of Evo-MD, we observe convergence at a ΔΔ*F_sensing_* that far exceeds the values we see for current state-of-the-art lipid packing defect sensors (e.g. HCV AH, ALPS and M2 MH).

With the highly efficient mechanical pathway method we present here, we can now perform a similar optimization for the lipid packing defect sensing problem, which would be unfeasible with previous methods, like TI, because of their computational expense. Fig. 7B illustrates the utility of Evo-MD in generating peptide sequences (with a fixed length of 24 amino acids) with highly diverse sensing free energies. We should emphasize that, in the current work, these results solely serve as a proof of principle and that details on the sequences of the resolved optima as well as their experimental validation will be published in a separate upcoming paper.

An important and fundamental question is whether the natural and nature-derived peptides that we studied in this work are optimal for lipid packing defect sensing. Intriguingly, the evolutionary convergence in Fig. 7B suggests that known curvature sensors in nature (like ALPS and HCV AH) are in fact far from optimal. We argue that the many evolutionary constraints imposed by nature’s complexity (e.g. solubility, protein-protein interactions, trafficking and many more) likely hinder the optimization of a single objective. This argument is in full agreement with the notion that curvature sensing in nature is a subtle balance between general membrane binding and specific curvature recognition. ^33^ As an addition to this, our results suggest that – since defect sensing also implies the active induction of leaflet strain – curvature sensors should conserve the structural integrity of the lipid membranes they adhere to. Consequently, the global optimum of a desired single objective such as defect or curvature sensing in our example may thus lie far outside the applicability domain of data-science based generative peptide models, i.e. generative models trained on large datasets of native sequences, because the training data is too distinct from the theoretical optima.

We argue that directed evolution approaches such as Evo-MD may offer a valuable benchmark platform for lipid force-fields: First, the primary aim of force-fields is to reproduce physicochemical driving forces – trends rather than absolutes – and therefore force-fields must at least reproduce the global physicochemical features of the optimized sequences. Second, Evo-MD yields sequences over the whole range in relative binding free energy by gradually maximizing the relevant chemical distinction between the peptides in a well-spaced manner.

Of course, a main limitation of this approach is that CG models such as Martini do not predict secondary and tertiary structure. However, we argue that secondary structure only marginally affects the CHAOS value, especially for peptides with a strong lipid packing sensing ability. We can already see this effect in the here-examined HCV AH peptide: Although HCV AH was originally assumed to fold into a straight *α*-helix,^38,48^ a recent study used a helix-turn-helix (HTH) tertiary structure for HCV AH^49^ as a starting point for their MD simulations, as predicted by PEP-FOLD.^50^ However, when we calculate CHAOS values for this HTH fold and compare it to the helical HCV AH, we find only minor differences between the two in terms of lipid packing sensing ability (see SI). These findings indicate that optimization of sensing is largely dominated by the peptide’s general amino acid composition (i.e. its overall hydrophobicity) rather than peptide structure, in line with similar findings in the context of membrane-binding antimicrobial peptides.^51^ This implies that one can effectively generate sequences with high CHAOS values and subsequently resolve the structure via atomistic simulations or – if the goal is direct comparison – restrain the structure in both the atomistic and CG simulation to, for example, an *α*-helix. In addition, the Evo-MD runs can be performed in conjunction with ‘on the fly’ structure prediction without severe loss of computational efficiency, since the MD simulation will remain the rate-limiting step.

## 4 Conclusions

We found that a peptide’s ability to sense lipid packing defects in biological membrane can be redefined as the peptide’s ability to reduce membrane tension in a leaflet under excess strain, or equivalently, the peptide’s ability to reduce the mechanical work required to stretch the membrane (leaflet). We have demonstrated that the calculation of such a reduction in mechanical work offers a highly efficient and accurate route for the estimation of relative membrane binding free energies. This resulting quantification of lipid packing sensing ability by the membrane peripheral protein can be expressed as the sensing free energy (ΔΔ*F_sensing_*), or the here-defined Characteristic Area of Sensing (CHAOS) parameter, which is independent on the choice of end-states. The method only requires knowledge of the individual pressure components over the course of the simulation trajectory to calculate the system’s surface tension, a quantity which virtually all molecular dynamics packages provide.

We used this novel end-state method to compare the performance of the new Martini 3 coarse-grained force-field with the previous Martini 2 for biorelevant lipid packing defect sensing peptides. We observed that Martini 3 most accurately reproduced the experimental trends, although the relative binding free energies were generally lower than the ones calculated from atomistic simulations. In contrast, Martini 2 severely overestimated peptide-membrane interactions and consequent packing defect sensing. Finally, since defect sensing implies both easing and active induction of leaflet strain, we hypothesize that lipid packing defect and curvature sensors in nature must be far form optimal since these proteins must largely conserve the structural integrity of the lipid membranes they adhere to. We argue that the extrema, i.e. the optimal sequences, particularly provide valuable benchmarks for force-field development and comparison because their distinct physicochemical signatures directly reflect how a force-field captures the physicochemical mechanisms of sensing.

## Acknowledgement

The authors thank Dr. Edgar Blokhuis and Max Makurat for the discussions regarding the CHAOS parameter definition. Jeroen Methorst is thanked for his insights in the Evo-MD results. The authors acknowledge the Dutch Research Organization NWO (Snellius@Surfsara) and the HLRN Göttingen/Berlin for the provided computational resources. This work was funded by the Deutsche Forschungsgemeinschaft (DFG, German Research Foundation) under Germany’s Excellence Strategy - EXC 2033 - 390677874 - RESOLV. The authors additionally thank the NWO Vidi scheme project number 723.016.005 (the Netherlands), and the DFG grant number RI2791/2-1 (Germany) for funding.

## Supporting Information Available

Derivation of the relation between relative strain and vesicle diameter. Peptide sequences and properties. The relation between sensing free energies in constant area and constant tension ensembles. CHAOS parameters compared to the area increase upon peptide adhesion. 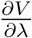-profiles for thermodynamic integration. Density plots of peptide-membrane binding in different force-fields. Insertion depths of peptide-membrane binding in different force-fields. Effect of HCV AH tertiary structure on CHAOS parameter.

