## supporting_information for "Efficient quantification of lipid packing defect sensing by amphipathic peptides; comparing Martini 2 & 3 with CHARMM36"

### 1 Derivation of the relation between relative strain and vesicle diameter

Membrane curvature can be modeled by the stretching and compression of two uncoupled elastic sheets. Consequently, when the outer monolayer is bent, the area of the neutral plane within the outer leaflet remains constant, whereas the region above this plane is stretched, and the region below this plane is compressed. Now, we define $d$ as the distance from the head groups to the neutral plane of the outer monolayer, i.e, approximately half the monolayer thickness, and $R$ as the distance from the vesicle center to this neutral plane (Fig. SI1). The relative strain $\epsilon$ (area increase) between the leaflet area at the position of the head groups $R + d$ and the neutral plane $R$ can be derived via the different spherical areas:

$$\epsilon = \frac{A - A_0}{A_0} = \frac{4\pi(R+d)^2 - 4\pi R^2}{4\pi R^2} = \frac{R^2 + 2dR + d^2}{R^2} - 1 = \frac{d^2}{R^2} + \frac{2d}{R} \tag{1}$$

Which can be rearranged to find the effective vesicle radius $R$ as a function of $\epsilon$:

$$R = \frac{d(1 + \sqrt{1+\epsilon})}{\epsilon} \tag{2}$$

This expression of R is equivalent to the result obtained by others,[1] with the difference that we base our approximation on the neutral plane of the outer monolayer rather than the bilayer when bending two uncoupled elastic sheets. Now, if we plug in the numbers for our system ($\epsilon = 0.165$ and $d \simeq 1$ nm), we find that this corresponds to an effective vesicle radius of $\approx$ 12.5 nm and, consequently, a diameter in the range of 25 nm. Though it should be noted that such a simple estimation remains rather approximate since it is based on the assumption of a uniform lateral stretching modulus.

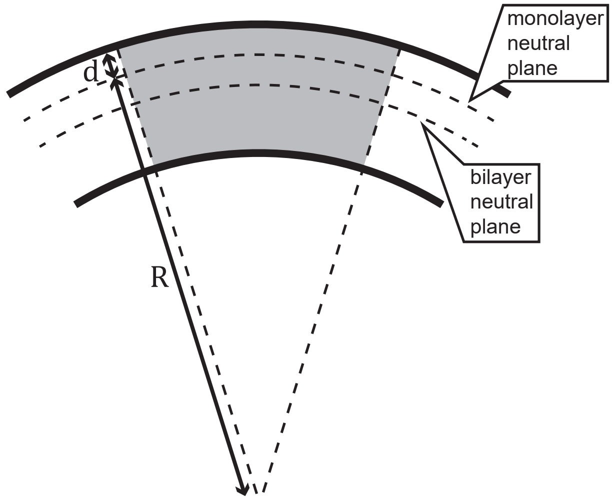


Figure SI1: Schematic depiction of a bent membrane, with the relevant distances $R$ and $d$.

### 2 Peptide sequences and properties

Table SI1: Sequences and physicochemical properties for the six peptides studied in this work. Mutated amino acids (in HCV NH and M2 MH/NH) with respect to the original peptides (HCV AH and M2 AH, respectively) are highlighted in gray. Hydrophobicities ($< H >$) and hydrophobic moments ($< \mu_H >$) were calculated for the full sequences by HeliQuest[2] (default settings).

| Peptide | Sequence | Length | Charge | $< H >$ | $< \mu_H >$ |
|---|---|---|---|---|---|
| ALPS | DDFLNSAMSSLYSGWSSFTTGASKFASAAKEGATK | 35 | 0 | 0.279 | 0.365 |
| HCV AH | SGSWLRDVWDWICTVLTDFKTWLQSKL | 27 | 0 | 0.686 | 0.465 |
| HCV NH | SGSWLRDDWDWECTVLTDDKTWLQSKL | 27 | -3 | 0.221 | 0.427 |
| M2 AH | RLFFKCIYRFFEHGLKRG | 18 | +4 | 0.519 | 0.474 |
| M2 MH | RKFFKKIYRFFRKLLKRL | 18 | +9 | 0.335 | 0.874 |
| M2 NH | RLAAKCAARFAEHGLKRG | 18 | +4 | 0.153 | 0.263 |

### 3 The relation between sensing free energies in constant area and constant tension ensembles

The stretching free energy at constant area ($\Delta F|_A$) can be extended to the free energy at constant tension ($\Delta G|_\sigma$) by using the Legendre transformation:

$$dG|_\sigma = \left(\frac{\partial G}{\partial A}\right)\Big|_A dA + \left(\frac{\partial G}{\partial \sigma}\right)\Big|_\sigma d\sigma = \sigma dA\big|_A + \Delta A d\sigma\big|_\sigma \tag{3}$$

Note that we only use $\Delta F$ to emphasize that the free energies are obtained within the constant area ensemble. However, both ensembles are performed at a constant pressure of $p_z = 1$ bar. When we integrate this between the tensionless state (labeled 0) and the high tension state (labeled 1), we find

$$\Delta G|_\sigma = \int_{A_0}^{A_1} \sigma dA\big|_A + \int_{\sigma_0}^{\sigma_1} \Delta A d\sigma\big|_\sigma = \underbrace{\frac{A_1 - A_0}{2}(\sigma_1 + \sigma_0)\Big|_A}_{\Delta F|_A} + \underbrace{(A_1' - A_1)\sigma_1\Big|_{\sigma_1}}_{\Delta G|_{\sigma_1}} - \underbrace{(A_0' - A_0)\sigma_0\Big|_{\sigma_0}}_{\Delta G|_{\sigma_0}} \tag{4}$$

for the system without a peptide adhered to the membrane surface. The first term ($\Delta F|_A$) is identical to the definition of eq. 6 in the main text. The two correction terms represent the constant tension contributions to the stretching free energies at high tension ($\Delta G|_{\sigma_1}$) and tensionless conditions ($\Delta G|_{\sigma_0}$). Similarly, for the system with an adhered peptide to the membrane surface we find:

$$\Delta G'|_{\sigma'} = \int_{A_0'}^{A_1'} \sigma' dA\big|_{A'} + \int_{\sigma_0'}^{\sigma_1'} \Delta A' d\sigma\big|_{\sigma'} = \underbrace{\frac{A_1' - A_0'}{2}(\sigma_1' + \sigma_0')\Big|_{A'}}_{\Delta F'|_{A'}} + \underbrace{(A_1' - A_1)\sigma_1'\Big|_{\sigma_1'}}_{\Delta G'|_{\sigma_1'}} - \underbrace{(A_0' - A_0)\sigma_0'\Big|_{\sigma_0'}}_{\Delta G'|_{\sigma_0'}} \tag{5}$$

As an example, we performed additional simulations for the ALPS peptide with surface-tension coupling to the surface tensions we found in the constant area runs for the four conditions ($\sigma_0$, $\sigma_0'$, $\sigma_1$ , and $\sigma_1'$). This showed that the correction terms for the tensionless membrane systems, $\Delta G|_{\sigma_0}$ and $\Delta G'|_{\sigma_0'}$, are indeed negligibly small ($0.12{\cdot}10^{-3}$ and $-6.29{\cdot}10^{-3}$ kJ mol$^{-1}$, respectively), because $\sigma_0$ and $\sigma_0'$ are – per definition – close to zero in the tensionless state. Also, the correction terms for the tensed states, $\Delta G|_{\sigma_1}$ and $\Delta G'|_{\sigma_1'}$, are nearly identical (0.697 and 0.650 kJ mol$^{-1}$, respectively), such that they almost entirely cancel out when calculating the sensing free energy ($\Delta\Delta G_{sensing}\big|_{\sigma,A} = \Delta G'|_{\sigma'} - \Delta G|_{\sigma}$).

Taken together, we find a corrected sensing free energy $\Delta\Delta G_{sensing}\big|_{\sigma,A}$ of $-12.077$ kJ mol$^{-1}$ for ALPS, which is merely 0.040 kJ mol$^{-1}$ smaller than the value we find when only simulating the systems at constant area ($\Delta\Delta F_{sensing}\big|_{A} = -12.037$ kJ mol$^{-1}$, see main text). From this, we conclude that the constant area simulations of the two end-states yield sufficiently reliably free energy values and do not require additional simulations of the constant tension ensembles.

### 4 CHAOS parameters compared to the area increase upon peptide adhesion

Table SI2: Calculated CHAOS parameters (eq. 8, Fig. 3D in main text) compared to the induced area upon peptide-membrane adhesion in an $NpT$ ensemble (tensionless conditions, average and standard error over 1 $\mu$s of simulation).

| Peptide | CHAOS parameter ($nm^2$) | Area induction in $NpT$ ($nm^2$) |
|---|---|---|
| ALPS | $0.611 \pm 0.029$ | $0.513 \pm 0.015$ |
| HCV AH | $0.691 \pm 0.032$ | $0.619 \pm 0.015$ |
| HCV NH | $0.034 \pm 0.007$ | $0.026 \pm 0.017$ |
| M2 AH | $0.408 \pm 0.022$ | $0.329 \pm 0.020$ |
| M2 MH | $0.566 \pm 0.028$ | $0.450 \pm 0.013$ |
| M2 NH | $0.202 \pm 0.012$ | $0.112 \pm 0.031$ |

### 5 $\frac{\partial V}{\partial \lambda}$ -profiles for thermodynamic integration

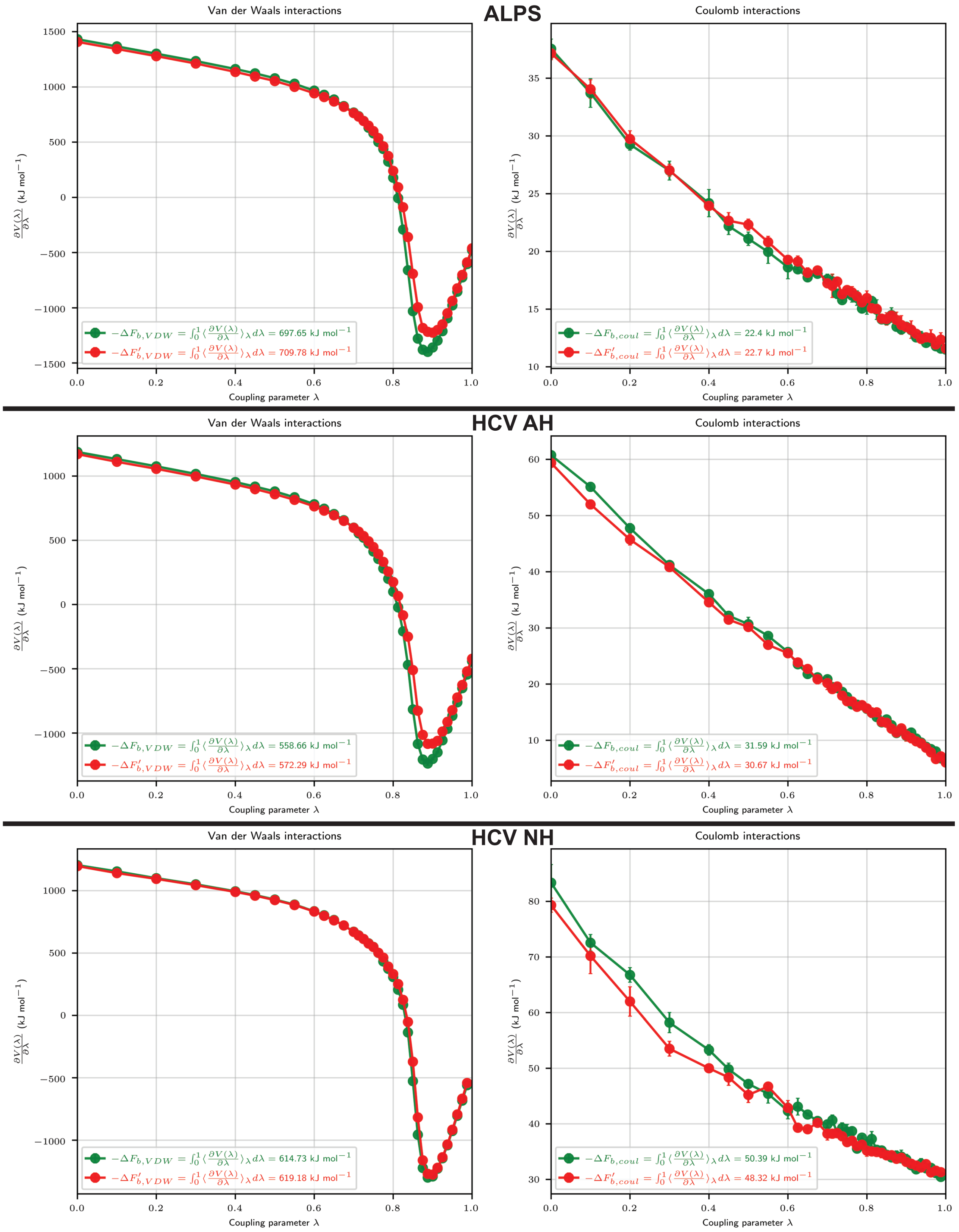

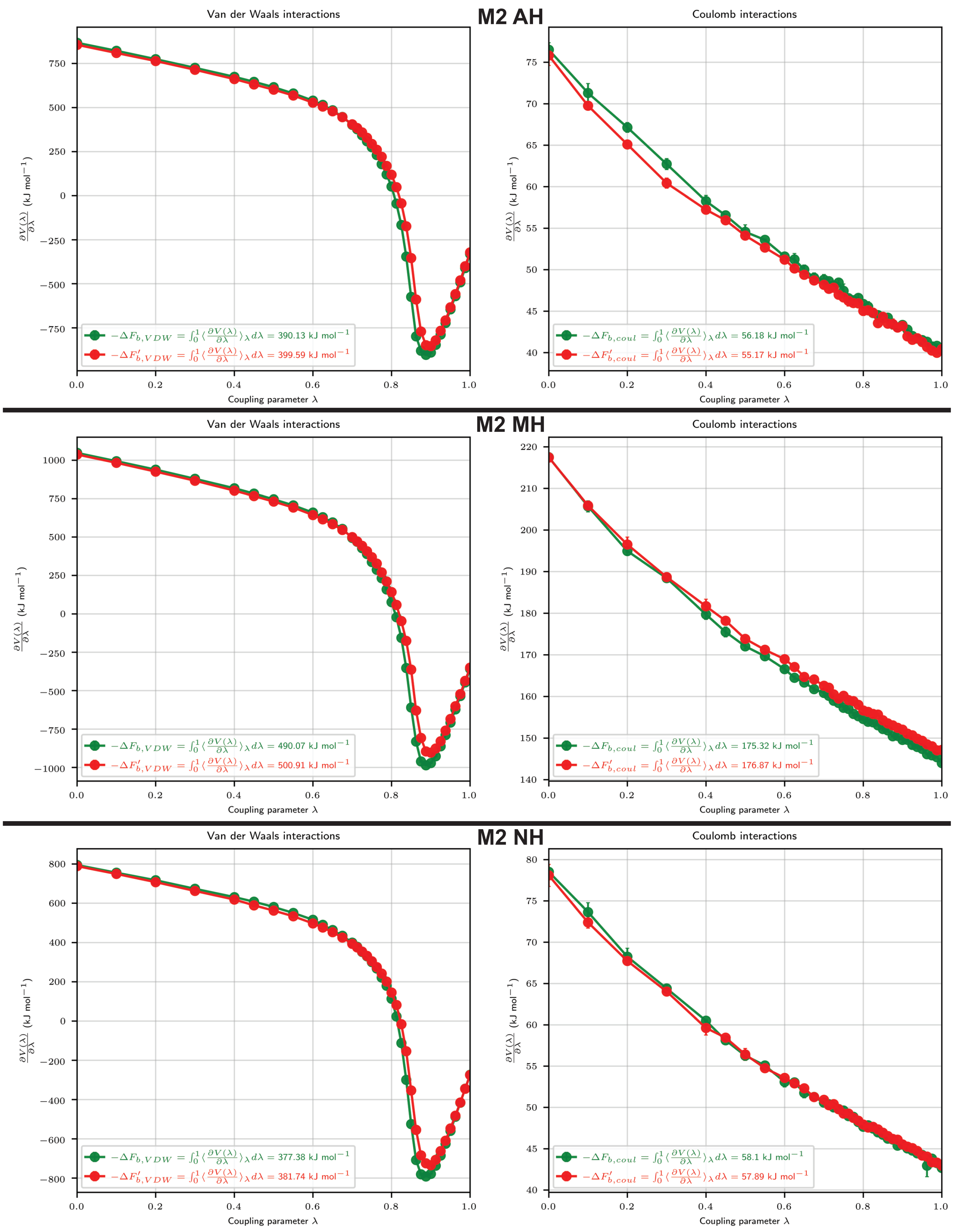


Figure SI2: The $\frac{\partial V}{\partial \lambda}$ -profiles for decoupling Van der Waals (left) and Coulomb (right) interactions for the tensionless system (green) and the tensed system (red). Integration of these curves yield free energy differences for the seperate interaction types (see legends). From these, we calculated $\Delta\Delta F_{sensing} = \Delta F'_b - \Delta F_b = (\Delta F'_{b,VDW} + \Delta F'_{b,coul}) - (\Delta F_{b,VDW} + \Delta F_{b,coul})$.

### 6 Density plots of peptide-membrane binding in different force-fields

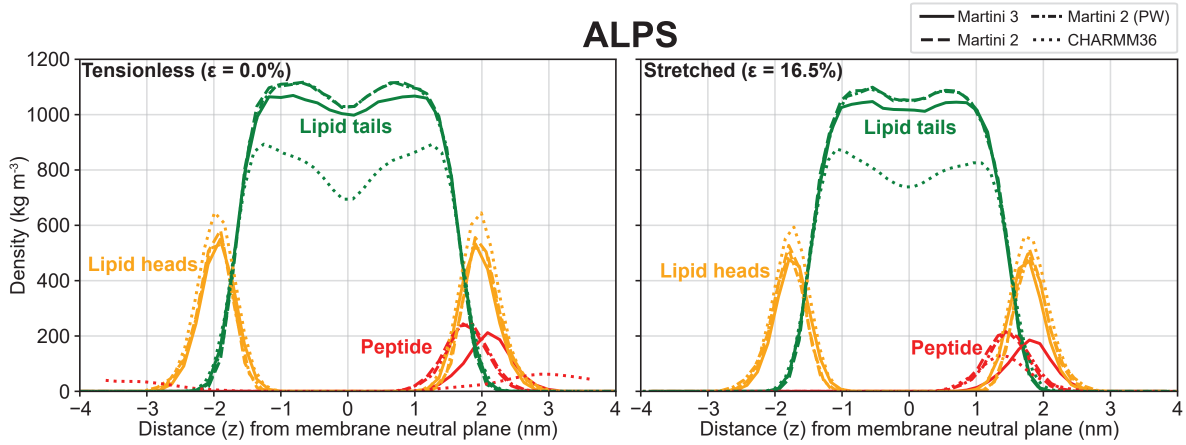


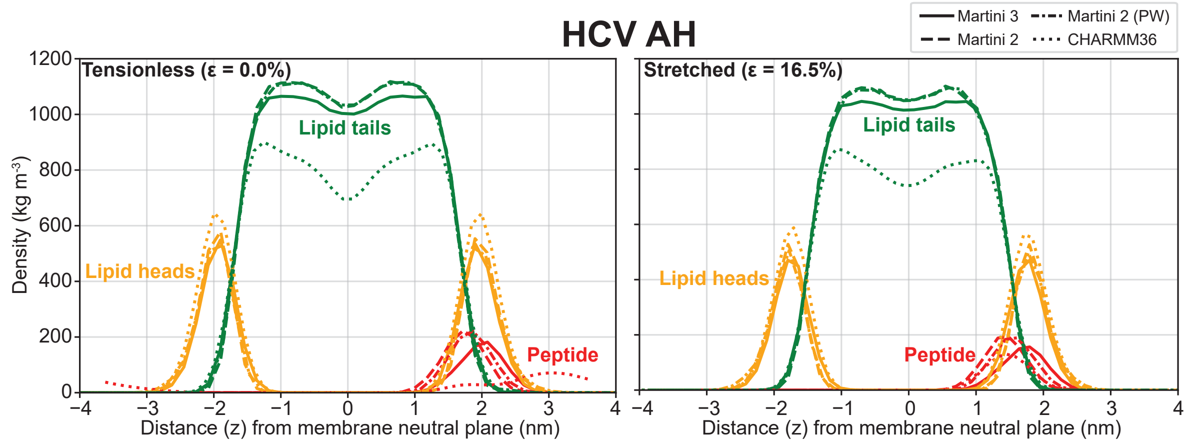


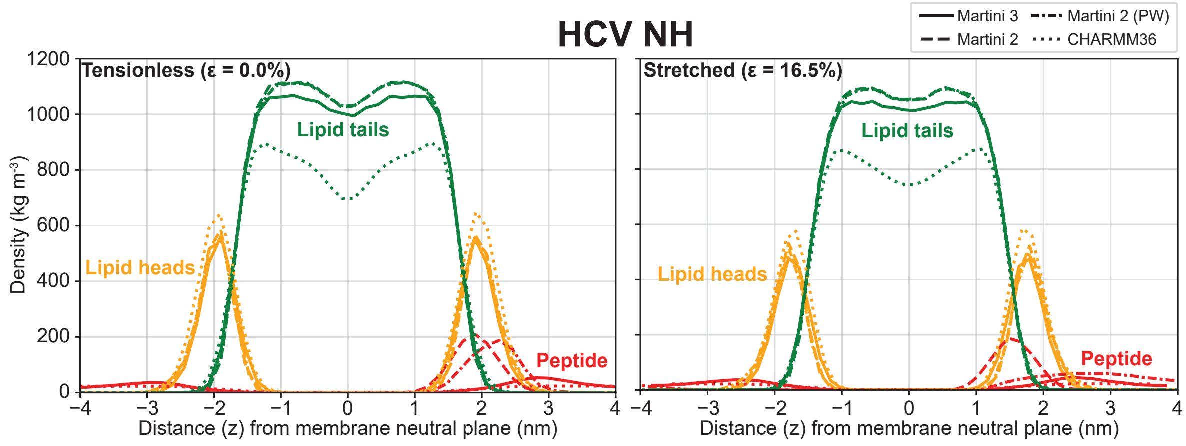

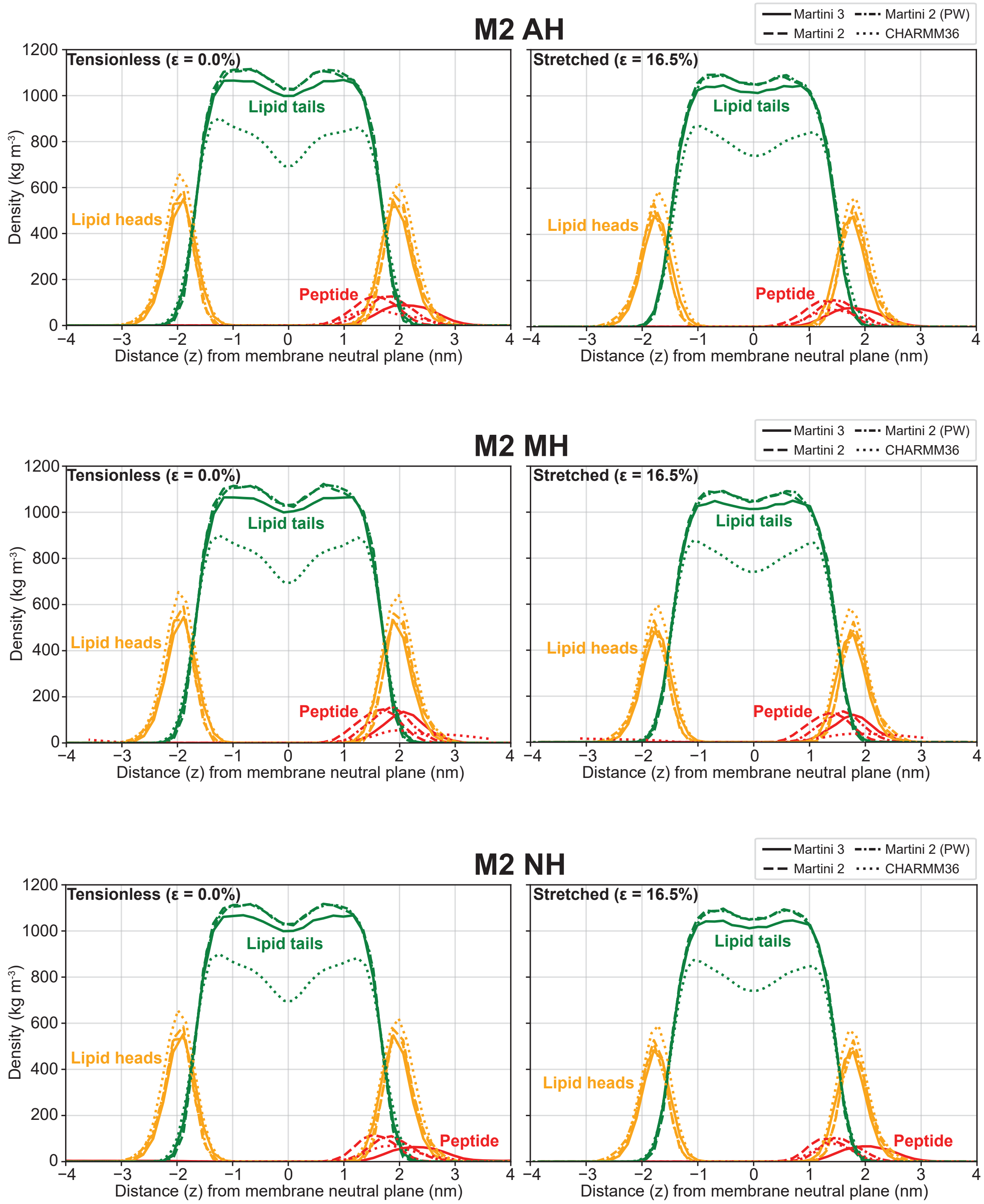


Figure SI3: Density plots calculated from MD trajectories of peptides on tensionless (left) and stretched (right) POPC membranes, using different force-fields. Phosphocholine groups were taken as the ‘lipid heads’. The alkyl tails and glycerol moieties are taken together as the ‘lipid tails’. Solvent densities are not shown for clarity.

### 7 Insertion depths of peptide-membrane binding in different force-fields

Table SI3: Distances (z) from the bilayer neutral plane (in nm) for the lipid head groups (monolayer thickness), and the individual peptides, as calculated from the density plots (Fig. SI3). Percentages represent the ratio between the peptide value and the lipid head group value.

| | **Martini 3** | | **Martini 2** | | **Martini 2 (PW)** | | **CHARMM36** | |
|---|---|---|---|---|---|---|---|---|
| | $\epsilon$=0.0% | $\epsilon$=16.5% | $\epsilon$=0.0% | $\epsilon$=16.5% | $\epsilon$=0.0% | $\epsilon$=16.5% | $\epsilon$=0.0% | $\epsilon$=16.5% |
| Lipid heads | 1.90 | 1.79 | 1.90 | 1.79 | 1.86 | 1.78 | 1.98 | 1.75 |
| ALPS | 2.09 (110%) | 1.79 (100%) | 1.74 (92%) | 1.44 (80%) | 1.78 (95%) | 1.48 (83%) | unbinding | 1.43 (82%) |
| HCV AH | 1.99 (105%) | 1.71 (96%) | 1.73 (91%) | 1.42 (79%) | 1.83 (98%) | 1.54 (87%) | unbinding | 1.45 (83%) |
| HCV NH | unbinding | unbinding | 1.90 (100%) | 1.55 (87%) | 2.18 (117%) | unbinding | unbinding | unbinding |
| M2 AH | 2.14 (113%) | 1.82 (102%) | 1.55 (82%) | 1.29 (72%) | 1.79 (96%) | 1.49 (84%) | 1.67 (84%) | 1.48 (85%) |
| M2 MH | 2.08 (109%) | 1.78 (100%) | 1.62 (85%) | 1.35 (75%) | 1.83 (98%) | 1.57 (88%) | unbinding | unbinding |
| M2 NH | 1.83 (96%) | 2.04 (114%) | 1.58 (83%) | 1.34 (75%) | 1.86 (100%) | 1.55 (87%) | 1.94 (99%) | 1.43 (79%) |

### 8 Effect of HCV AH tertiary structure on CHAOS parameter

We compared the peptide structure predictions for HCV AH of PEP-FOLD[3] with the recent (state-of-the-art) AlphaFold2[4] and found that the former predicts a helix-turn-helix (HTH) folding (Fig. SI4A) and the latter predicts an $\alpha$-helix (Fig. SI4B). This already indicates that the folding of HCV AH is not trivially defined, likely very dynamic and dependent on the environment (in aqueous solvent versus membrane-bound). We simulated both conformations in Martini 3 found that there is very little difference between the CHAOS parameters for HCV AH in an $\alpha$-helical ($0.691 \pm 0.032$ nm$^2$) or a HTH configuration ($0.636 \pm 0.023$ nm$^2$).

To illustrate the dynamics of HCV AH in the presence of a (tensionless) membrane, we analyzed the end-to-end distance between the the peptide's termini (S1 and L27 backbone/$C_\alpha$ atoms) in our atomistic and CG trajectories, starting from an $\alpha$-helix (Fig. SI4C). In line with the known limitations of the Martini model considering protein folding, we observed that the CG model retains the initial $\alpha$-helical configuration (long end-to-end distance). In contrast, the atomistic model dynamically transitions to a partly unfolded HTH conformation with a shorter end-to-end distance (Fig. SI4C), and low affinity for the tensionless membrane (consistent with previous findings[1]). It is important to note that our current data do not provide fully conclusive answers in this regard, since both peptide folding and membrane binding are processes that have timescales of multiple microseconds and therefore usually require enhanced sampling methods to capture properly, which is beyond the scope of the current work.

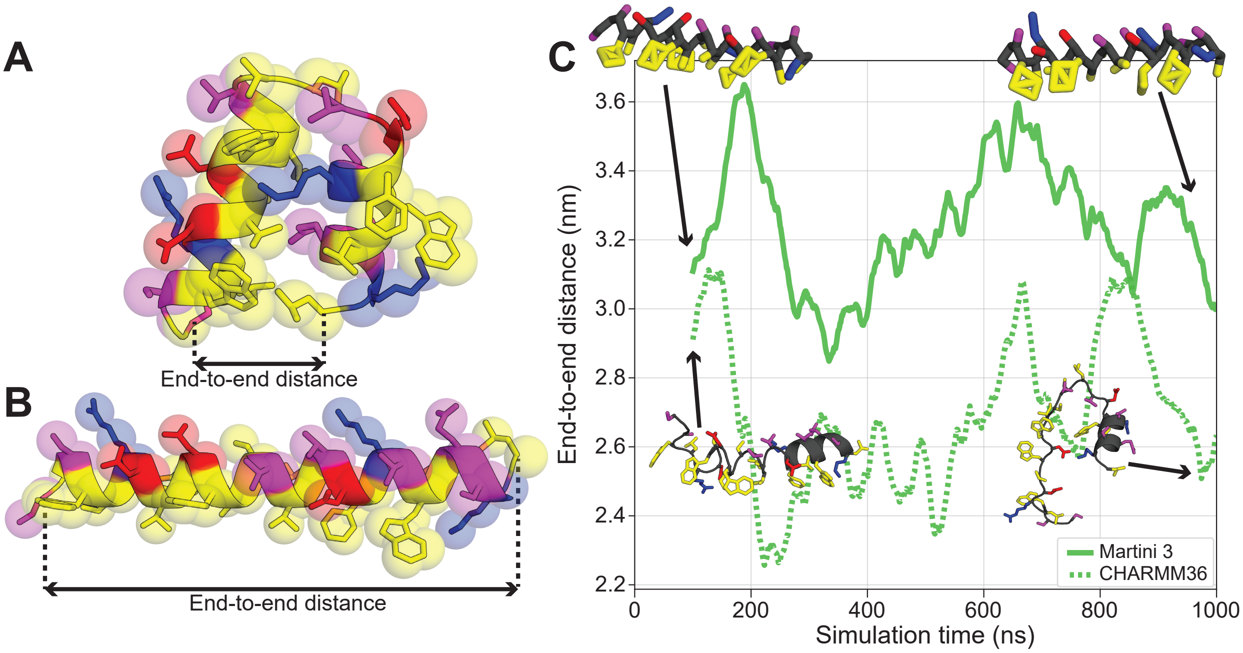


Figure SI4: Predicted atomistic structures for HCV AH by PEP-FOLD[3] (A) and AlphaFold2[4] (B). Transparent spheres represent the Martini 3 CG beads. C) The end-to-end distance between the atomistic and CG peptide termini (100 ns moving average) in the presence of a tensionless membrane. Insets: atomistic and CG snapshots of the peptide structures (lipids and solvent not shown for clarity).
